# Modelling red blood cell optical trapping by machine learning improved geometrical optics calculations

**DOI:** 10.1101/2023.02.25.530023

**Authors:** R. Tognato, D. Bronte Ciriza, O. M. Maragò, P. H. Jones

## Abstract

Optically trapping red blood cells allows to explore their biophysical properties, which are affected in many diseases. However, because of their nonspherical shape, the numerical calculation of the optical forces is slow, limiting the range of situations that can be explored. Here we train a neural network that improves both the accuracy and the speed of the calculation and we employ it to simulate the motion of a red blood cell under different beam configurations. We found that by fixing two beams and controlling the position of a third, it is possible to control the tilting of the cell. We anticipate this work to be a promising approach to study the trapping of complex shaped and inhomogeneous biological materials, where the possible photodamage imposes restrictions in the beam power.

## Introduction

In 1970 Arthur Ashkin first demonstrated how to manipulate and confine microscopic particles suspended in water through radiation pressure [3]. Following the first demonstration of optical trapping, Ashkin and collaborators developed the single-beam gradient trap, today known as optical tweezers (OT) [5, 40]. The basic principles of OT utilise the fact that light carries momentum which can be harvested to manipulate microscopic particles in solution. In its conventional and simplest set-up, OT focus a collimated Gaussian laser beam to a diffraction-limited spot where it can trap microparticles. Soon after the first demonstration of OT, Ashkin et al. employed them to manipulate biological particles like bacteria and erythrocytes without causing damage [4, 6].

In humans, erythrocytes, or red blood cells (RBCs), are anucleated cells responsible for the oxygen delivery to tissues and organs. Mature and healthy RBCs have a biconcave disk shape that minimizes the membrane bending energy. Typically, RBCs have diameter of 6-8 *µ*m, a peripheral thickest portion of 2-3 *µ*m, and a central dimple 0.8-2 *µ*m thick [18]. The excess surface area and membrane elasticity render the cell elastic and permits the RBC to pass through the microvasculature by deforming [30]. Alterations in the RBCs’ membrane elasticity are implicated in severe disfunctions of the microcirculation (e.g., capillaries can be entirely clogged, triggering tissue necrosis or organ damage and failure) [1]. The RBCs’ alteration can be genetically inherited [16], a consequence of a pathogen infection [41], a metabolic disorder [2], or due to radiation treatment [25]. Very recently, it has also been correlated to SARS-Cov2 infection [31].

In the last decades, OT have been widely applied in RBC research to investigate biochemical and biophysical properties of both healthy and unhealthy RBC via single- or multi-beam OT [7]. In these studies, researchers have adopted two main approaches to trapping: the indirect trapping, where handles as silica or polystyrene microspheres are used to manipulate the RBC [36], and the direct trapping, where the light beam directly traps the RBC [6]. Regardless of the mechanism used for trapping, the nature of biological samples makes them particularly susceptible to photodamage. To minimise this, infrared light in the second biological window (wavelength around 1064 nm) is generally preferred for the experiments [8].

As the cell is significantly larger than the incident wavelength, the geometrical optics approximation (GO) models properly the beam cell interaction [21, 33, 38]. GO assumes that the beam can be discretized in a series of rays that carry a fraction of the total momentum and by calculating and summing up the scattering of all rays, it is possible to compute the total force applied. However, even though GO simplifies considerably the theoretical treatment compared to a full wave optical approach [27], an accurate calculation requires consideration of a large number of rays, with an associated high cost in computational time. Simulating the Brownian dynamics requires repetition of the force calculation at each time step sequentially, which becomes prohibitively slow if the force calculation is not optimized [10]. The fact that the calculation is sequential and that the shape is complex prevents the use of conventional approaches to speed up the calculation (e.g., parallelization and interpolation based approaches).

Machine learning, and in particular neural networks (NN), are emerging across a variety of research fields as a powerful technique to solve challenging problems. Backed by their ability to learn from previous examples in order to make new predictions, NN are contributing to biology [29], food sensing control [17], and even to containment of epidemics [35]. In fact, NN have recently been demonstrated as an useful technique to increase both the speed [32] and the accuracy [9] of optical forces calculations when compared to GO, allowing the study of more complex systems through Brownian dynamics simulations. While these previous works consider spheres [32] and ellipsoids [9], there is no evident reason to remain constrained to these relatively simple shapes. Indeed, the computation time saving that could be achieved by using NN for force calculation of particles with more complex shapes makes this a particularly attractive application.

In this work, we train a NN to enhance the speed and accuracy of the optical force calculation for RBC. This permits a numerical exploration of the Brownian dynamics of a RBC, potentially allowing to study in a more complete manner different trapping configurations. More efficient trapping configurations employ less laser power and therefore reduce the risk of photo damaging the trapped cells.

## 1 Methods

### 1.1 Model and geometrical optics calculations

In our model we consider a Gaussian beam propagating along the opposite direction of the force of gravity (+*z* direction). The wavelength (1.064 *µ*m) is selected to match the vast majority of experiments and in agreement with previous works on trapping RBCs [2, 42]. The OT parameters are the ones of a typical OT experiment (beam power 5 mW, numerical aperture 1.3).

The RBC is assumed to be in its healthy biconcave disk conformation, and the parameters describing the shape of the RBC are those reported by Evans et al. [18]. A radius (*r*) of 3.91 *µ*m, a central dimple with a thickness (*t*_*min*_) of 0.81 *µ*m, and a thickest portion, located at 2.76 *µ*m from the axis of symmetry, with a thickness (*t*_*max*_) of 2.52 *µ*m. According to the Evans-Fung model, the thickness (*Z*) of a section of the RBC reads:

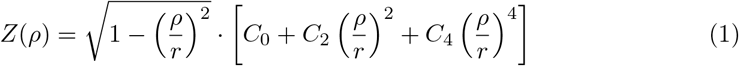

where *ρ* is the radial distance from the axis of symmetry, *r* is the cell radius and *C*_0_, *C*_2_ and *C*_4_ (0.81, 7.83, -4.39, respectively) are numerical values related to the observable parameters that describe the cell morphology [39].

As RBCs are significantly larger than the wavelength of the incident light, the optical forces acting on them can be calculated with GO. We perform this calculation with the specialized software OTGO [11]. For biological samples, such as RBCs, that have a low refractive index contrast with the typical suspending medium, the fraction of power that is reflected after a scattering event is very low (*<* 0.001) [22], therefore in our ray tracing calculations, only the first two scattering (refraction) events are considered.

### 1.2 Diffusion tensor

The erratic motion of a particle trapped in liquid in an OT set up is influenced by the fluid’s resistance, by the thermal noise, and by the external deterministic forces exerted by the OT [19, 27]. For non-spherical objects, a single scalar diffusion coefficient is not enough to describe the statistics of the random motion. It is necessary to use a 6 × 6 diffusion tensor (**D**), which depends on the particle shape and orientation [27]:

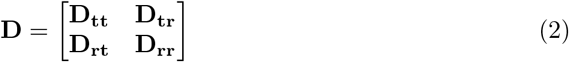

where **D**_**tt**_, **D**_**rr**_ and **D**_**rt**_ = **D**_**tr**_^T^ are 3 × 3 blocks and the subscripts ‘r’ and ‘t’ refer to the particle’s rotational and translational degrees of freedom, respectively.

Although an analytical expression for (**D**) exists for simple shapes like spheres, ellipsoids or cylinders [24], the RBC morphology is more complex and requires numerical methods for its determination. Here, we used the bead model technique developed by De La Torre et al., exploiting the widely used software winHYDRO++ [15, 20]. In the bead model, a series of spheres are used to approximate the size and the total volume of the RBC. From the bead model, winHYDRO++ calculates the 6×6 tensor (**Ξ**) encoding the hydrodynamic resistance of the non-spherical particle. We then obtain the diffusion tensor **D** via the generalised Einstein relationship [14]:

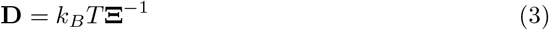

where *k*_*B*_ is the Boltzmann constant, and *T* is the temperature of the system.

In the present study, the bead model is constructed in a strict sense, filling the volume of the RBC considering only spheres of equal sizes. In this case, the centre of diffusion of the particle coincides with the centre of mass of the particle, and the numerical output for the diffusion tensor reads:

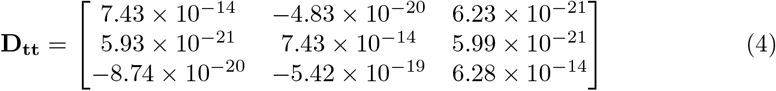

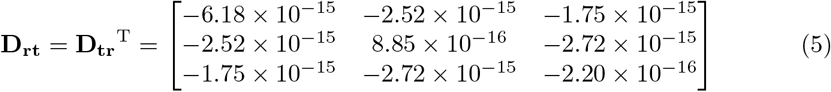

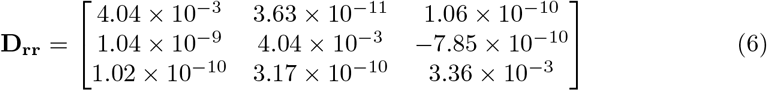

where the units are 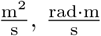, and 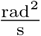 respectively. Notice that the diagonal terms of **D**_**tt**_ and **D**_**rr**_ that indicate the diffusion coefficient along a specific direction (i.e., x,y,z) and a specific axis (i.e., x,y,z) are several orders of magnitude larger than the off diagonal terms highlighting the shape-induced directional dynamics typical of non-spherical particles [23].

### 1.3 Particle dynamics simulation

The simulation of the dynamics of the RBC is based on the works of M. X. Fernandes et al. [19], and described in reference [27]. Two reference frames are defined: a particle reference frame Σ_p_, which has an origin that coincides with the particle’s centre of mass (CM) and the centre of diffusion (CD), and a laboratory reference frame Σ_*l*_ that is centred at (0,0,0) and which axes are oriented along 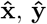, and 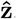, Fig. 1-a. At time *t*, the RBC’s CD is located at **r**_CD_(*t*) = [*x*_CD_(*t*), *y*_CD_(*t*), *z*_CD_(*t*)]. The cell orientation can be described by the angles *α*_1_(*t*), *β*_1_(*t*) and *γ*_1_(*t*) defined with respect to the particle unit vector 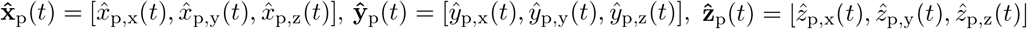. **D** is obtained in the particle reference frame, that is centred at **r**_CD_(*t*) and the axes are oriented along 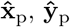 and 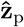

**Figure 1.**
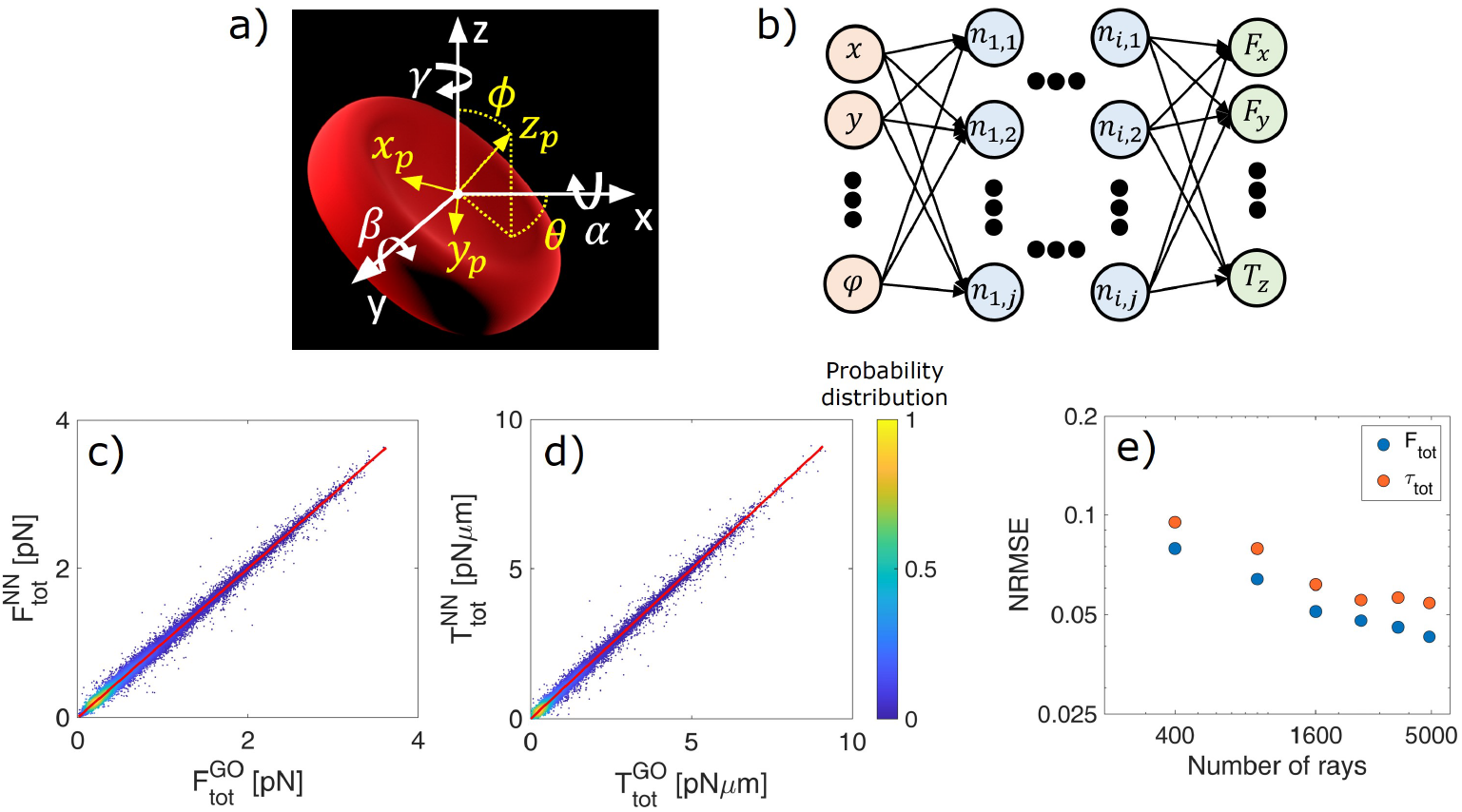
a) Definition of particle (yellow) and laboratory (white) reference frames and rotation angles (*α, β, γ*) of the cell around the laboratory reference frame; b) schematic depiction of the neural network. The input layer contains six neurons describing the cell position and orientation, and the output layer has six neurons describing the components of force and torque acting on the cell. In between are seven hidden layers (*i* = 7), each of them with 256 neurons (*j* = 256). c-d) Density plots comparing the magnitude of the total force 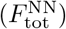 and torque 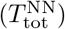 predicted with NN with those calculated with the GO method 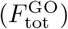 and 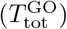. Regression lines are shown in red. e) Log-Log plot of the normalised root mean squared error (NRMSE) between 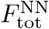 and 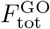, and 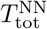 and 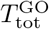 as a function of the number of rays used in the GO calculation. For each data point, the NN employed remains the same (trained with 4 *×* 10^2^ rays).

To simulate the free diffusion of an arbitrarily shaped particle from time *t* to the time step *t* + ∆*t*, initially one has to calculate the increment of the particle position and orientation in Σ_p_(*t*):

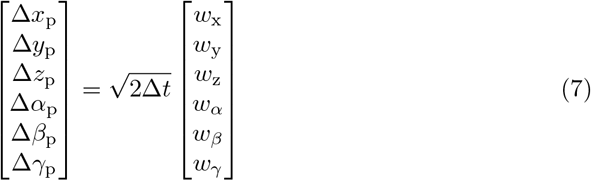

where [*w*_x_, *w*_y_, *w*_z_, *w*_*α*_, *w*_*β*_, *w*_*γ*_]^T^ are white noise terms, random numbers obtained from a multivariate normal distribution with zero mean and covariance **D**. Successively, the increments of the particle position calculated in Σ_p_ has to be transformed to Σ_*l*_. This is given by the transformation matrix:

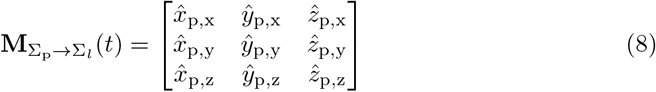

Therefore, the finite difference equation to update the particle position in Σ_*l*_ is:

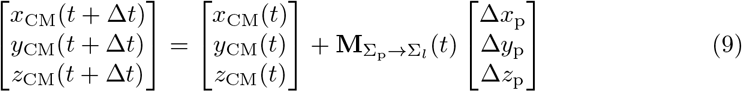

Once the new particle position is calculated, one has to update the particle orientation from Σ_p_(*t*) to Σ_*l*_(*t*), which is effectively a rotation of the particle unit vectors. This rotation, for small angles, is expressed in Σ_p_ by the rotation matrix:

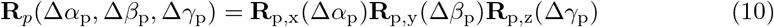

where

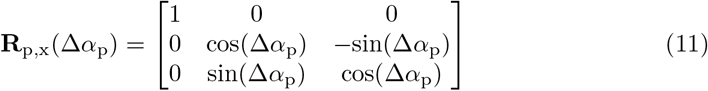

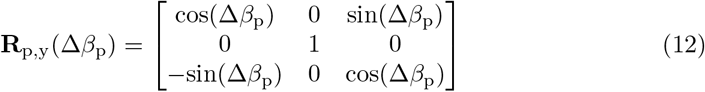

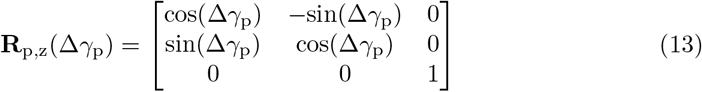

Transforming this rotation matrix to Σ_*l*_, we obtain the unit vectors representing the orientation of the particle at the end of the time step:

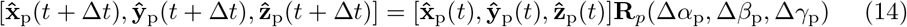

As the last step, the rotation matrix has to be updated:

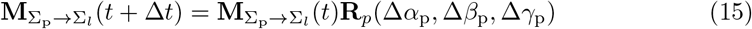

However, in the current situation, we must also account for the optical forces (**F**) and torques (**T**) exerted by the optical trap on the centre of mass of the RBC. Therefore, taking into account **F** and **T**, the increments of the particle orientation and position in Σ_p_ are:

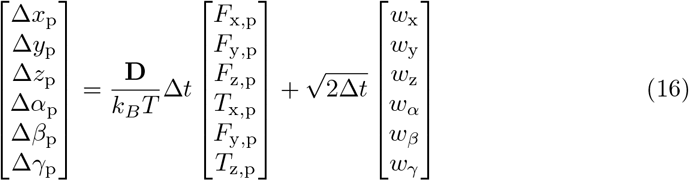

which then need to be transformed back into Σ_*l*_. However, **F** and **T** are calculated in Σ_*l*_, and therefore they must be transformed to Σ_p_ via the matrix 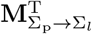.

The finite difference scheme is combined with the NN or with the GO code to calculate the optical forces and torques acting on the RBC and to simulate the Brownian dynamics of the optically trapped particle. We estimate the time step, ∆*t*, for the Brownian motion simulation from the trap stiffness reported by Tognato et al. [38], and from the diffusion properties of a healthy RBC obtained here. The typical time scale on which the restoring force acts is given by 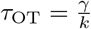, while the momentum relaxation time is given by 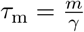. To assure numerical stability, ∆*t* must fall in between these two characteristic time scales (*τ*_OT_ *≫* ∆ *t ≫τ*_m_) [27]. From the diffusion tensor **D**, one can extract the diffusion properties of the RBC along a specific direction (D_i_), then through the fluctuation-dissipation theorem one can obtain *γ*_*i*_. For example, for the x-direction one obtains 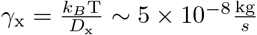. Therefore, considering a trap stiffness in this direction 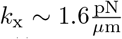, one obtains *τ*_OT_ *∼* 3 *×* 10^*−*2^s. On the other hand, given a mass of *∼*1 *×* 10^*−*11^kg for a typical healthy RBC, one obtains *τ*_m_*∼* 4 *×*10^*−*4^s.A similar estimation can be made for the other directions, and contemplating the magnitude of the other terms in **D**, a time step ∆*t* = 0.001s is adequate to assure the numerical stability [27].

### 1.4 Neural network architecture and training

The neural network (NN) architecture is composed of one input layer with 6 neurons representing position and orientation of the cell (*x, y, z*, cos(*θ*), sin(*θ*), *ϕ*), one output layer with 6 neurons representing the force and torque components (*F*_x_,*F*_y_,*F*_z_,*T*_x_,*T*_y_,*T*_z_), and 7 hidden layers in between with 256 neurons each, Fig. 1-b. While Σ_p_ encodes the orientation of the particle by using 3 angles, because of symmetry, the orientation of the RBC can be completely defined by the polar angle *ϕ* and the azimuthal angle *θ*, as shown in Fig. 1-a.

The training data consists of 4 *×* 10^6^ different points in the 5D space of parameters (*x,y,z,θ,ϕ*). 90% of these data points are used as a training data set while the remaining 10% is kept as a testing data set to evaluate the accuracy of the NN. The training data are generated via GO calculations made in OTGO [11]. The cell is placed in uniformly distributed positions in a cube of side 8*µ*m centred at the origin of the Cartesian coordinates system (i.e. *−*4*µ*m *≤*x*≤ −*4*µ*m, *≤* 4*µ*m*≤* y *≤* 4*µ*m and *−*4*µ*m*≤* z *≤*4*µ*m). Simultaneously, to account for the possible different orientation of the RBC within the trap, the cell is uniformly and randomly oriented in an interval for *− π ≤ θ ≤ π* and 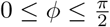. The training data are generated for the simplest case of a single-beam OT with the geometrical focus centered at (0, 0, 0) and a beam power of 5mW.

The NN is trained in Python using Keras (version 2.2.4-tf) [13]. The training of the NN is divided into 5 different steps. The data pre-processing and the model definition, which are done only once, and the loading of the data, the training step, and the evaluation of the performance, that are carried out iteratively. The training data, generated as previously described, contains data in different units and scales. While the position scale is in the order of *∼*1 *×*10^*−*6^m, the forces are on the range of*∼* 1 *×* 10^*−*12^N, and the torques are around *∼*1 *×* 10^*−*18^N · m. To achieve an efficient training of the NN, we need to apply a pre-processing step where the variables must be rescaled around unity and *θ*, that ranges from *−π* to *π*, is expressed in terms of sines and cosines to avoid inconsistencies around 2*π*. Shuffling the data and dividing them into a validating and training set is the final step of the pre-processing. In our case, the training data set contains 5.4 *×* 10^6^ points while 6 *×* 10^5^ points are reserved for the testing data set. In this work, we employ fully connected NNs where each neuron is activated by a sigmoidal function. Defining the model implies choosing the number of layers and the number of neurons per layer. Among the explored architectures, the one consisting of 7 hidden layers provides the best results (in terms of accuracy, training time, and speed).

The iterative part of the training starts by loading the training data and applying the training step where the NN weights are optimised to minimise the loss function. We use the mean squared error as the loss function and the Keras implementation of the Adam optimiser [13]. Once the training dataset is fully explored, the difference between the NN calculation and the validating dataset (defined as the mean square difference) is computed. The iterative step is repeated until this difference reaches its minimum value and we consider that the model is fully trained. The training of the NN is done in a GPU type NVIDIA GeForce RTX 2060 with 16 GB of memory. The processor of the computer is an Intel Core i7-10700, and it has 16 GB of RAM.

## 2 Results

### 2.1 Single beam Optical Tweezers

To evaluate the effectiveness of our approach, we start by testing the ability of the NN to predict the forces and torques acting on an RBC in a single beam OT (SBOT). We compare the NN predictions (trained with data generated using 4 *×* 10^2^ rays) and the*×* GO calculations considering 4 times more rays (1.6 *×*10^3^ rays) at 1*×* 10^5^ random positions and orientations. The 2D density plots shown in Fig. 1-c and d illustrate the agreement between the NN and GO in predicting the optical forces (regression coefficient 0.998, *R*^2^ = 0.996) and torques (regression coefficient 0.999, *R*^2^ = 0.996), respectively. We further demonstrate the accuracy of the NN by comparing our NN (trained with data generated with 4 *×*10^2^ rays) with the GO calculation (considering a greater number of rays). Fig. 1-f shows the normalised root mean squared error (NRMSE) between the predictions of the NN trained with 4 *×*10^2^ rays and the GO calculations with different numbers of rays (up to 5*×* 10^3^ rays). The NRMSE decreases as the number of rays increases. The forces and torques calculated with 5 *×* 10^3^ rays result more similar to the NN output than to the forces obtained with a total of 4 *×* 10^2^ rays, meaning that the NN is able to increase the accuracy of the force and torque prediction, even for an object with such a complex shape.

### 2.2 Double beam Optical Tweezers

Since the NN is trained for a SBOT, one may think it can only predict the optical forces and torques for a SBOT. However, the NN can be used multiple times to simulate multi-beam optical tweezers. In fact, the NN can predict the forces generated by a single beam on different locations on the cell, and the total force acting on the centre of mass of the cell may then be calculated as the vector sum of each contribution. Here we consider a double-beam optical tweezers (DBOT) where the two beams’ geometric foci are positioned 5.06*µ*m apart along the *x*-axis, similar to the experiments conducted by Agrawal et al. [2], Fig. 2-a.

**Figure 2.**
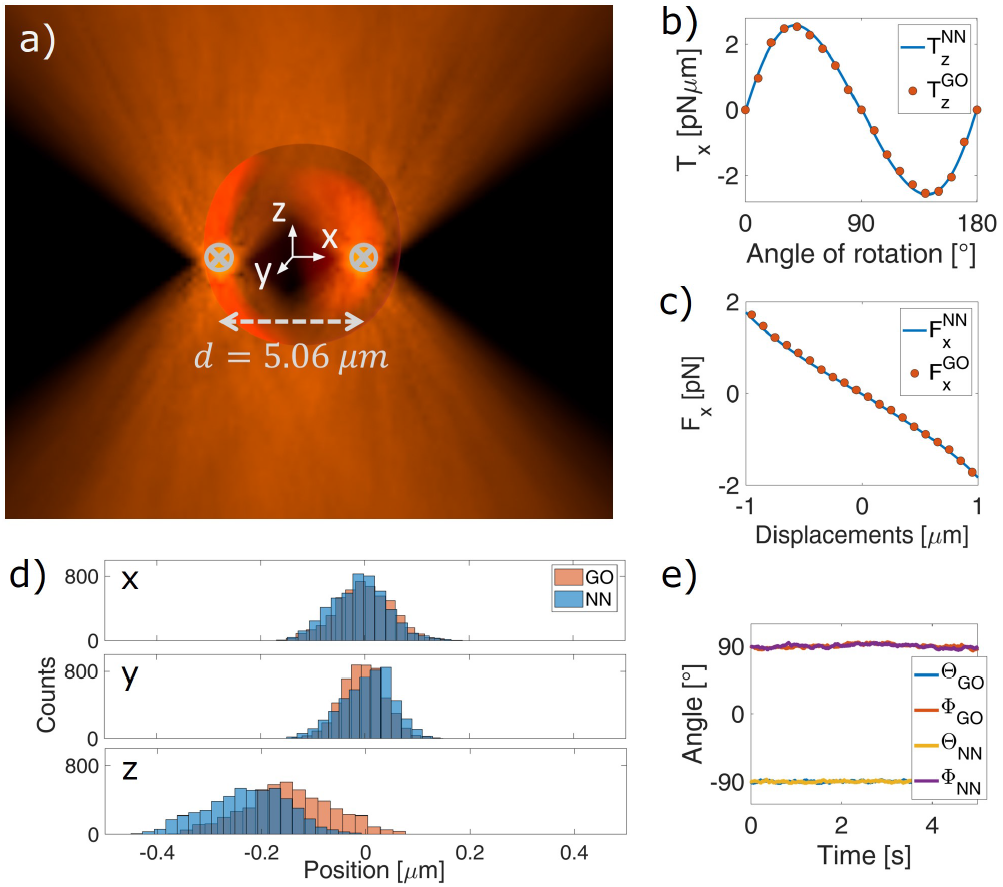
a) Schematic depiction of an RBC trapped by a double-beam OT. b-c) Comparison between the GO calculation and the NN prediction for the b) torque-rotation curve for rotation around the x-axis and c) force-displacement curve along the x-direction. (d) Comparison of the probability distribution obtained with the GO calculation and with the NN prediction for a RBC in a DBOT. (e) Cell orientations in the numerical simulation for both GO and NN.

Fig. 2-(b-c) shows *T*_x_(*α*) and *F*_x_(*x*) calculated with GO and predicted with the NN for a cell in its folded configuration (i.e., cell major axis parallel to the optical axis) trapped in a DBOT. In both cases, the NN predictions (solid line) agree well with the GO method (dots), demonstrating the possibility to use the NN for multi-beam optical traps. We therefore conclude that this approach can be extended to predict forces and torques generated by a three- and four-beam OT, situations in which the GO calculation is considerably slower given the very large number of light rays required.

We now investigate the cell’s dynamics within a DBOT using both NN and GO to compute the optical forces. The simulation of the Brownian dynamics follows the strategy explained in the Methods section (Particle dynamics simulation) where now, the force and torque considered is the sum of the contributions of each of the beams. Fig. 2-(d) shows the probability distribution of the centre of mass of the cell for a total simulation time of 5s, while Fig. 2-(e) shows the orientation of the cell with respect to the fixed reference frame as a function of the simulation time. It is important mentioning that in the current configuration a rotation around the y-axis (*β*) would be a rotation around the cell axis of symmetry and therefore completely irrelevant. By extracting the average values for each degree of freedom, it is possible to compare the final equilibrium configuration obtained with the NN and with GO. Indeed, the average values obtained with the predictions of the NN and GO methods agree well and are also in good agreement with previously reported values [38], Table 1.

**Table 1.** Equilibrium position and orientation for a RBC in a double-beam OT as found with geometrical optics (GO) and with neural networks (NN). For each parameter we report the average and the standard deviation.

|  | GO | NN |
| --- | --- | --- |
| $x_{2,eq}(\mu\text{m})$ | $0.01 \pm 0.05$ | $0.01 \pm 0.05$ |
| $y_{2,eq}(\mu\text{m})$ | $0.00 \pm 0.05$ | $0.00 \pm 0.04$ |
| $z_{2,eq}(\mu\text{m})$ | $-0.20 \pm 0.08$ | $-0.18 \pm 0.08$ |
| $\phi_{2,eq}(^{\circ})$ | $90.65 \pm 2.11$ | $90.27 \pm 1.44$ |
| $\theta_{2,eq}(^{\circ})$ | $-90.45 \pm 1.06$ | $-90.09 \pm 0.92$ |

Moreover, the biggest advantage of using the NN for numerical simulations is a consistent decrease in the simulation time required to achieve the same precision (the NN is two orders of magnitude faster). Since the NN shows a higher computational efficiency, hereafter, we make use of the NN prediction to simulate the Brownian dynamics of an optically trapped RBC.

We therefore move to extract quantitative information on the trap constants. Initially we analyse the hydrodynamics of the RBC, since non-spherical particle could have an intrinsic roto-translation coupling due to their peculiar shape [23]. In our case, the diffusion tensor **D** does not show any strong roto-translation coupling; therefore, we do not expect to find any strong correlation in the cell’s motion intrinsically due to the RBC’s hydrodynamics. Still, optically trapped non-spherical particles could show roto-translation coupling in their motion as previously observed by others. In this framework, the normalised auto-correlation function (ACF) has been successfully used to extract quantitative information about the trapping constants [28, 34].

We first evaluate the spatial ACFs (*C*_xx_ (*τ*), *C*_yy_ (*τ*), *C*_zz_ (*τ*)) of the particle centre of mass trajectories. *C*_xx_ (*τ*) and *C*_zz_ (*τ*) decay as a single exponential with characteristic decay frequencies *ω*_x_ = 28 s^*−*1^ and *ω*_z_ = 6.4 s^*−*1^. Contrariwise, *C*_yy_ (*τ*) is well fitted with a double exponential with characteristic frequencies *ω*_*y*,1_ = 42 s^*−*1^ and *ω*_*y*,2_ = 2.7 s^*−*1^, Fig. 3-a. We associate the fast decay rate to the translation, while the slower decay can be related to rotation around the *x*-axis (*α*) induced by a motion along the y-direction. The normalised cross-correlation function between *α* and *y* Fig. 3-b further confirm a roto-translation coupling (amplitude -0.368) [26]. Fig. 3-c shows a density plot of the rotation around the *x*-axis (*α*) as function of the motion along the *y*-direction. Here it can be seen a moderate negative correlation which suggests that the RBC rotates as it moves away from *y*_eq,2_, and undergoes to an “oscillating” motion about the equilibrium configuration where it is stably confined. To better comprehend this correlation we simulate *F*_y_ (*α*) (Fig. 3-d) and *τ*_x_ (*y*) (Fig. 3-e) which undoubtedly shows the coupling between the motion along *y* and *α*. Actually, the cell in its “folded” position (i.e. *α* = 90^*◦*^) is constantly subjected to a force along the *y*-direction that moves the particle away from *y*_eq,2_ which in turns induces a rotation around the *x*-direction. On the other hand, as extensively described by Tognato et al., the transverse forces and torques components confine the cell in its “folded” configuration [38]. The overall consequence of these stable and unstable equilibria is a “circulating motion” of the cell within the optical trap about the equilibrium configuration. This would suggest that the coupling is intrinsically due to particle shape and to the optical trap rather than to the hydrodynamic of the particle.

**Figure 3.**
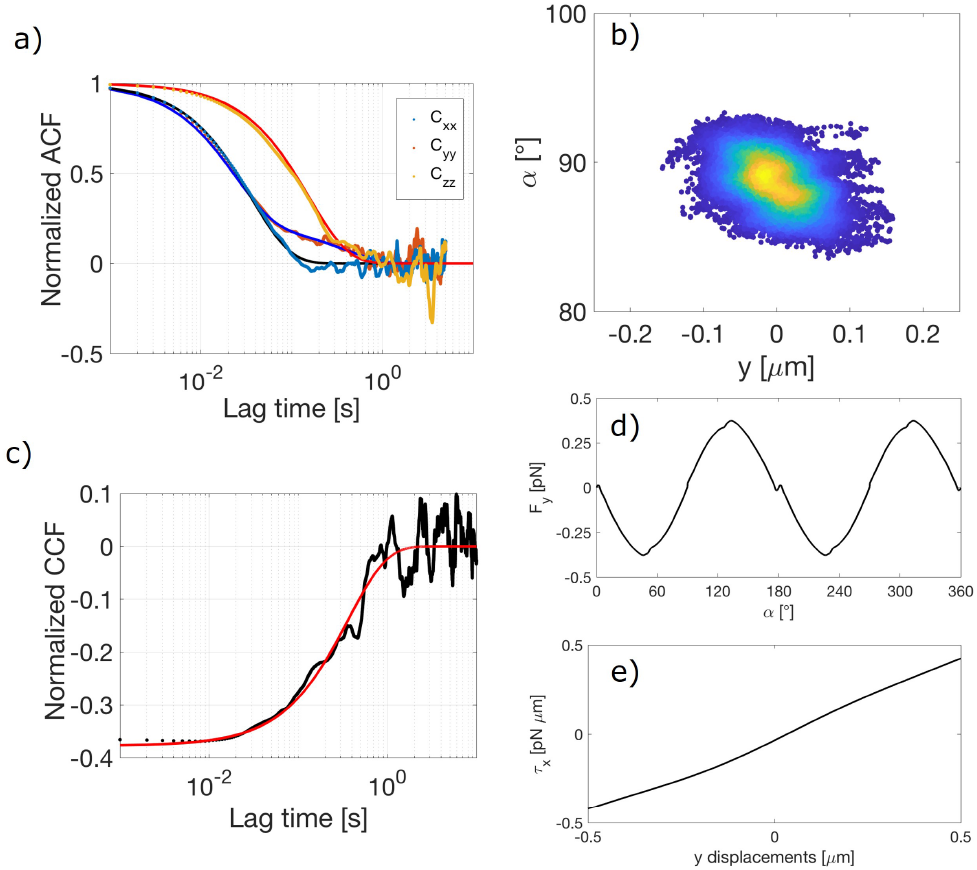
a) Translational autocorrelation function. The solid lines are exponential fits. *C*_*xx*_ (*t*), *C*_*zz*_ (*t*), decay as single exponential while *C*_*yy*_ (*t*) as double exponential. b) *y− α* correlation shown as density plot. c) Normalised cross-correlation function between the rotation around the x-axis (*α*) and the y-displacement (red line exponential fit). Both *F*_*y*_ (*α*) d) and *T*_*x*_ (*y*) e) reveal unstable equilibrium when the cell is tilted of 90° around the x-axis (i.e. RBC in its folded position)

Lastly, we extract average values and the standard deviations for the force constants 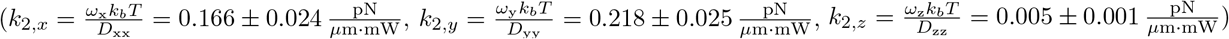. These values are in excellent agreement with a previously reported work [38]. Similarly to the translational motion, we calculate *C*_*αα*_ (*τ*) and *C*_*γγ*_ (*τ*), Fig. S1. *C*_*αα*_ (*τ*) and *C*_*γγ*_ (*τ*) decay as a single exponential and the respective trap constant are: 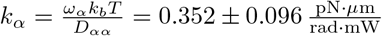 and 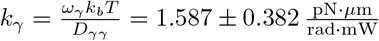. We do not analyse the dynamics around *β* since the cell is not confined about this axis.

### 2.3 Triple beam Optical Tweezers

As previously suggested, one of the greatest advantage of using a NN instead of GO is the significant lowered computation time, especially when a very high number of light rays is needed (e.g. a triple- or four beams optical tweezer). Now, we exploit this feature to investigate the equilibrium orientation and position of a RBC with a reconfigurable triple-beam OT.

If directly trapped, a healthy biconcave RBC can assume two different and alternative orientations within the optical trap depending on the number of beams used for trapping [7, 38]. In a double-beam OT, the major axis of a RBC is parallel to the optical axis and the beam foci are contained in the cell, known as “folded” configuration [2]. On the contrary, if three or four beams arranged in symmetric configurations are used (i.e. beams foci on the vertex of equilateral triangle or a square), the major axis of the cell is confined to be orthogonal to the optical axis (i.e. *α* = 0^*◦*^), configuration referred to as ‘flat’ configuration [37]. Here, we sought for alternative (and intermediate) RBC equilibrium configurations in respect to the well-known “folded” and “flat” ones.

We consider a trap configuration that is intermediate to those traps able to trap the cell in its “folded” or “flat” configuration. We consider a triple-beam optical tweezers (TBOT) composed by three identical and tightly focused Gaussian laser beams. Two beams are always arranged along the *x*-axis in a diametrically opposite location on the thickest portion of the cell (white crosses in Fig. 4-a). A third beam (yellow cross in Fig. 4-a) can be translated over the thickest portion of the cell and is used to counteract *T*_x_ generated by the two fixed beams. For simplicity, henceforth, the position of the moving beam is described by a polar co-ordinates system in the *x − y* plane. Its location is defined by a single angle (*ξ*), and the distance from the origin is fixed and equal to the radius of the thickest portion of the cell (2.76*µ*m), Fig. 4-a.

**Figure 4.**
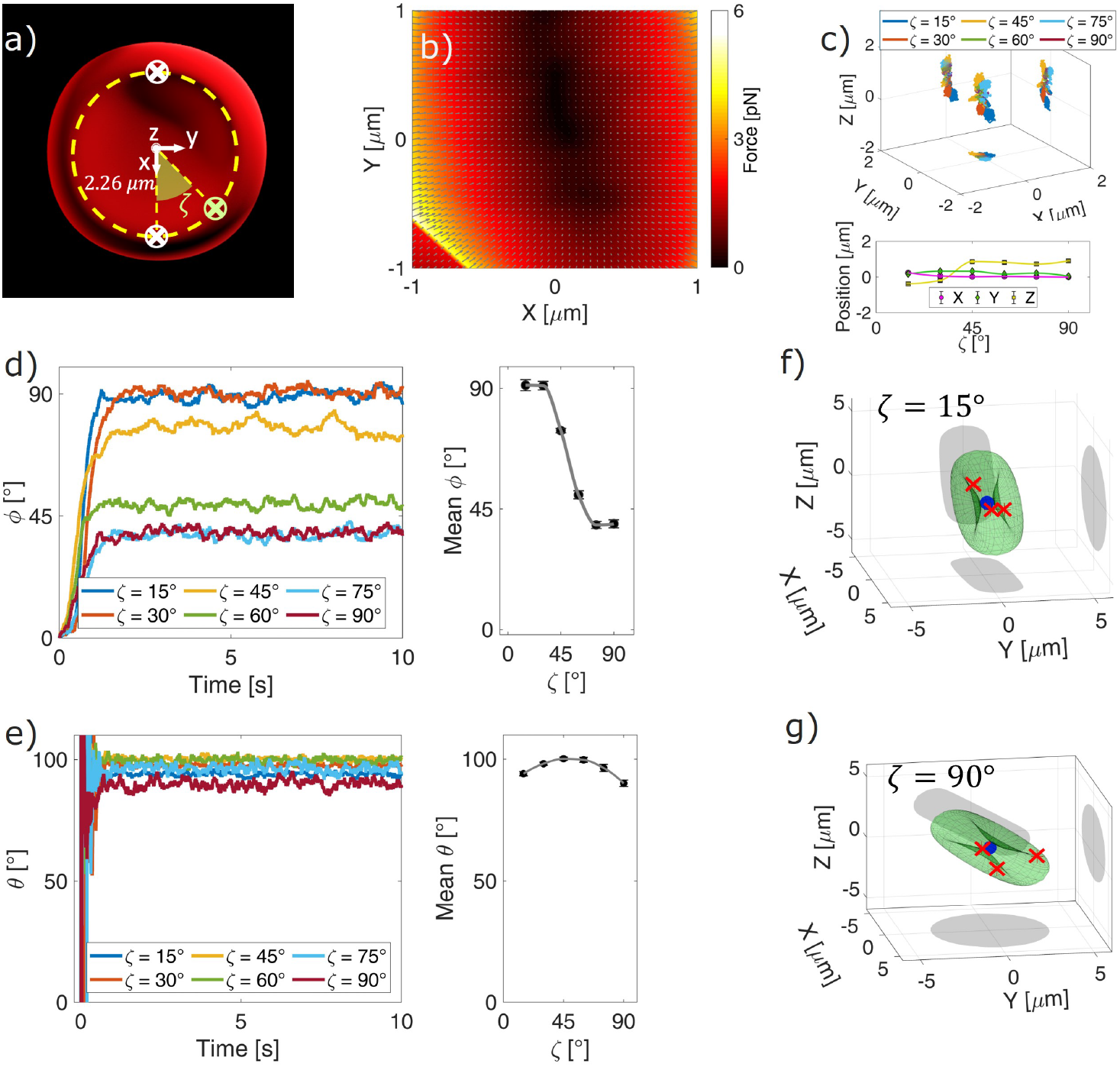
a) Schematic depiction of the triple-beam optical trap and the polar co-ordinates system used to identify the position of the moving beam. (b) Force-field for an RBC on the x-y plane for *ξ* = 45^*◦*^. The colour code indicates the total force acting on the x-y plane, while the grey arrows indicate the direction of the force. (c) Three-dimensional trajectories of the cell centre of mass over a simulation time of 10 s for different *ξ*, and the average values for the last second of simulation. d) Polar (*ϕ*) and e) azimuthal (*θ*) orientation of the RBC as a function of the simulation time. Average orientations are measured over the last second of the simulation. The error bar represents the standard deviation. f,g) Final equilibrium configuration for a RBC in the reconfigurable triple beam optical trap for *ξ* = 15^*◦*^ and 90^*◦*^ respectively. The blue dot indicates the center of mass of the RBC while the red stars indicates the beams’ foci.

Next, we proceed with the identification of the positional and translational equilibria. As a first step in our investigation, we simulate a force-field acting on the cell for *ξ* = 45^*◦*^ to appreciate the effect of the potential landscape on the RBC. In this simulation, the cell is in its “flat” configuration and located at *z* = 0. It can be seen that the light pattern creates a very complex force-field (Fig. 4-b). Non-negligible optical forces act simultaneously along the *x−* and *y*-direction for every location of the cell. The complexity of the force-field makes extremely difficult the identification of the equilibrium positions (i.e., point in space where a specific force component vanishes with negative slope). This process would requires several reiterations for every degree of freedom rendering the process labour intensive. However, noting that if a particle is subjected to an optical potential it falls into the equilibrium position/orientation, it would be possible to identify the equilibrium configuration studying its dynamics as suggested by Cao et al [12]. From symmetry arguments, the effect of different locations of beam 1 can be understood restricting *ξ* in the interval [0^*◦*^, 90^*◦*^] as schematically depicted in Fig. 4-a. Moreover, since we are looking for alternative equilibrium configurations (or to a transition from a “flat-like” to “folded-like” configuration), it is also rationale to disregard every position where two beams are too close to each other (i.e. *ξ <* 15^*◦*^), which should induce a “folded” configuration. Thus, the position of beam 1 can be restricted to 15^*◦*^*≤ ξ ≤* 90^*◦*^. To evaluate the effect of the reconfigurable optical trap *ξ* is sampled every 15^*◦*^, and for each *ξ*, the Brownian dynamic is simulated for a 10 s trajectory starting from a RBC positioned in its ‘flat’ configuration (*θ* = 0^*◦*^ and *ϕ* = 0^*◦*^) centred at (0, 0, 0). The simulation finishes once the cell equilibrates around a stable position and orientation. The final position and orientation are then given as the average position and orientation with the standard deviation of the last second of the simulation. Fig. 4-c shows the 3D trajectories of the RBC’s CM obtained from the simulations carried out for different *ξ*. Here, while *x* and *y* equilibrium positions remain close to the origin for different angles, the equilibrium in *z* does depend on *ξ*. In particular, for *ξ <* 30^*◦*^, *z*_*eq*_ *<* 0*µ*m and for *ξ >* 45^*◦*^, *z*_*eq*_ *>* 0*µ*m, Fig. 4-c. We anticipate that for *ξ* 30^*◦*^, the cell is in its ‘folded’ configuration, Fig 4-d and Fig. 4-f. This is due to a combination of the light intensity distribution and the cell configuration within the trap. In fact, when the cell is in its “folded” position, the cell’s major axes are parallel to the direction of propagation of the light beam. In this condition, more highly converging “light rays” strike the biggest faces of the RBC. This increases significantly the gradient force (*F*_*g*_). Simultaneously, while in folded position, the scattering force (*F*_*s*_) decreases appreciably because of the smaller geometrical cross-section of the cell. However, if *ξ* increases, this effect is less pronounced since the light rays strike the cell less symmetrically, and for *ξ* = 30^*◦*^, *z*_*eq*_ *∼ −*0.2*µ*m. Conversely, for *ξ ≥* 45^*◦*^ a net shift in the axial position is evidenced (*z*_*eq*_ *∼*0.8*µ*m), and this is due to a sequential shifting from the “folded-like” configuration to a “flat-like” configuration, Fig. 4-c (ii) and Fig. 4-d (i). Much more interesting is the analysis of the rotational equilibrium. In Fig. 4-d are shown the polar orientation (*ϕ*) of the cell as a function of the simulation time for different locations of the moving beam (i.e. various *ξ*). It is evident that *ξ* strongly influences the final polar orientation of the cell, Fig. 4-d. In particular, as beam 1 approaches beam 2, the cell tilts more until it reaches the “folded” configuration (i.e. *ϕ* = 90^*◦*^) for *ξ* = 30^*◦*^. Analysing the final orientation of the cell in more detail, it is possible to discriminate between three different regions. When the two beams are close to each other, *ξ ≤* 30^*◦*^, the cell is in the “folded” configuration. If 30^*◦*^*≤ ξ ≤* 75^*◦*^, the RBC’s tilting seems to vary linearly with *ξ*, from a “folded-like” configuration to “flat-like” configuration. The last region is for *ξ ≤* 75^*◦*^, where the cell tilting cannot be decreased further, Fig. 4-d. It is also interesting to note the minor effect that *ξ* has on *θ*. In particular, for *ξ* = 45^*◦*^ is possible to obtain the highest cell’s tilting around thez-axis, Fig. 4-e. For every other *ξ*, the tilting of the RBC around the z-direction decreases towards *θ* = 90^*◦*^.

## 3 Conclusions

The biomechanical properties of RBC are affected in different diseases and OT have been successfully used to trap RBC and to extract information about the membrane properties. While studying the optical forces applied on a trapped RBC can be done, the computation is extremely time consuming. In this work we have demonstrated that by using NN, one can significantly increase the speed of calculation without compromising the accuracy. This enhancement of the force calculations has allowed us to explore systems that were almost impossible to tackle with the conventional method for optical force calculation. In particular, we have focused on the analysis of the dynamics of trapped RBC with multiple beams. This can potentially allow determination of the best trapping configuration and to minimize the incident laser power and therefore reduce the risk of photodamaging the trapped cells.

## 5 Acknowledgments

The authors would like to thank Prof. Giovanni Volpe and Dr. Agnese Callegari (University of Gothenburg) for fruitful discussions on machine learning enhanced geometrical optics calculations.

